# Non-Parametric Mixture Modelling and its Application to Disease Progression Modelling

**DOI:** 10.1101/297978

**Authors:** Nicholas C. Firth, Neil P. Oxtoby, Silvia Primativo, Emilie Brotherhood, Alexandra L. Young, Keir X.X. Yong, Sebastian J. Crutch, Daniel C. Alexander

## Abstract

Dementia is characterised by its progressive degeneration of cognitive abilities. In research cohorts, detailed neuropsychological test batteries are often administered to better understand how cognition changes over time. Understanding cognitive changes in dementia is of great importance, particularly in determining how structural changes in the brain may affect cognition and in facilitating earlier detection of symptomatic changes. Disease progression models are often applied to these data to understand how a disease changes over time from cross-sectional data or to disease trajectories from large numbers of individuals. Previous disease progression models used to build longitudinal models from cross-sectional data have focused on brain imaging data; however, these models are not directly applicable to cognitive data. Here we use the novel, non-parametric, Kernel Density Estimation Mixture Modelling (KDEMM) approach and demonstrate accurate modelling of the progression of cognitive test data. We found that using KDEMM resulted in more accurate models of disease progression in simulated data compared to Gaussian Mixture Models (GMMs) for the majority of parameters used to simulate the data. When comparing KDEMM and GMM to cognitive data collected in different Alzheimers Disease subtypes, we found the KDEMM resulted in a model much more in line with clinical phenotype. We anticipate that the KDEMM will be used to integrate cognitive test data, and other non-normally distributed datasets into complex disease progression models.

## I. Introduction

Currently over half a million people in the UK have a diagnosis of dementia, this is projected to grow to one million by 2025 and two million by 2050 (*Diagnoses in the UK*). As there is currently no disease modifying therapeutic in most dementias, there is urgent need to gain better understanding of neurodegenerative disease to prevent eventually mitigate personal, societal and economic costs. It is estimated that measurable changes occur up to twenty years before a diagnosis of typical (memory led) Alzheimer’s disease (tAD) is given (Villemagne et al. 2013). This makes dementia particularly challenging to study because by the time of diagnosis it is difficult to discern early physiological or cognitive changes, here termed biomarkers, from more recent changes. Currently a large array of biomarkers are used to identify and characterise disease progression in the study of dementia. Examples include brain imaging (Slattery et al. 2017), cognitive (Pavisic et al. 2017), demographic (Singh-Manoux et al. 2017), genetic (Premi et al. 2017) and fluidic (Weston et al. 2015). A better understanding of the dynamics of these biomarkers will aid identification of which biological processes can be interrupted, ultimately to prevent further neurodegeneration and cognitive decline. Studying the dynamics of biomarkers *in vivo* is challenging given the prohibitive cost of preclinical studies, requiring enrolment of because it is prohibitively expensive to enrol large numbers of participants into studies which follow people forover long periods of time in order to observe pre-symptomatic changes. As such, disease progression models, which can reconstruct long-term pictures of disease from relatively short-term longitudinal, or even entirely cross-sectional, data sets. are used to infer biomarker dynamics in populations. Common varieties of model include: hypothetical (Jack et al. 2010), machine learning-based (Young et al. 2013), regression-based (Bilgel et al. 2016), Event-Based Model (Fonteijn et al. 2011), continuous trajectory (Villemagne et al. 2013; Donohue et al. 2014) and spatiotemporal (Lorenzi et al. 2015). With the exception of hypothetical models, these methods offer the potential to understand long-term biomarker dynamics on a common time frame from realistic data sets. The development of these methods was motivated in large part by the availability of large imaging data sets and their application to date has focussed mostly on imaging data.

In contrast to brain imaging, assessments of complex cognitive datasets have for the most part relied on traditional statistical approaches rather than data-driven methods. Optimising measures of detecting cognitive change is important not only for improving disease characterisation and prognosis in affected individuals, but also detecting and predicting change in asymptomatic at risk (sporadic) and presymptomatic (genetic) individuals (Dubois et al. 2016). Further optimising these assessments is important as neuropsychological differences between cognitively healthy subjects who will develop a dementia and those who will not can be observed 10 to 17 years before the diagnosis of dementia (Amieva et al. 2014). The evaluation of longitudinal change within and across different cognitive domains presents a number of specific challenges. First, performance across cognitive tasks is not independent. General factors (e.g. disease severity) and collateral deficits (e.g. visuoperceptual problems limiting performance on a face based memory test) can influence testing across many cognitive domains. Second, in many cases, cognitive profiles across tasks, both cross-sectionally and longitudinally, are described qualitatively, because test properties and normative samples differ across tasks. Third, the psychometric shape of tests differs markedly. Some tests yield relatively linear score distributions among healthy control participants because they contain graded difficulty items; other tests yield skewed score distributions owing to an excess of very easy or very difficult items. These psychometric properties influence the likelihood of clinical populations showing ceiling or floor effects at any given point in their disease progression. Fourth, practice effects mask longitudinal change. Practice effects across serial assessments (e.g. test familiarity, reduced anxiety) may conceal evidence of cognitive instability or decline (Machulda et al. 2017).

The key challenge faced in disease progression modelling is aligning multiple participants to an average model. It is not trivial to align participants temporally because: onset occurs at different ages; disease progression occurs at different rates (Buckley et al. 2016); and it is not practical to monitor large cohorts of presymptomatic participants to observe conversion, in particular when studying rarer forms of dementia. Two approaches to modelling these data can therefore be used; either some common time-frame can be inferred by aligning participants using a model (Donohue et al. 2014; Lorenzi et al. 2017), or temporal data can be disregarded and data can be aligned using some measure of disease severity. For a full review of different disease progression models see Oxtoby, *et al* 2017. The key challenge to overcome when aligning participants with a binary diagnosis is that some biomarkers only become noticeably affected late in the disease time course, where as others may be so sensitive that participants presumed to be healthy controls have impaired performance relative to normative samples in these measurements.

Here we develop a new kind of Event-Based Model (EBM) designed specifically to work with cognitive data. The key innovation is to use Kernel Density Estimation (KDE) to provide a non-parametric model of the cognitive score distribution models that underpin the EBM. We compare the new KDE method with a ubiquitous parametric mixture modelling technique, Gaussian Mixture Models (GMM). GMMs and KDE mixture models are compared using a goodness of fit metric and by their ability to recreate event-sequences and disease stages from synthetic data. We also show a use case of our non-parametric EBM, which yields the first comparison of cognitive deterioration in Posterior Cortical Atrophy (PCA) and typical Alzheimers disease (tAD). PCA is a clinico-radiological syndrome characterized by progressive decline in visual processing and other posterior cognitive functions, relatively intact memory and language in the early stages, and atrophy of posterior brain regions (Benson, Davis, and Snyder 1988; Crutch et al. 2017). PCA is most commonly caused by Alzheimers disease, with greater amyloid plaque and/or neurofibrillary tangle distribution in the posterior cortices than individuals with an amnestic presentation (Tang-Wai et al. 2004; Hof et al. 1997). Detailed longitudinal studies of cognitive change in PCA are currently lacking.

## II. Methods

In this section, we will describe the EBM and how mixture modelling is a major component of the model fitting. We will then discuss mixture modelling in detail, firstly the current methods used and then we will detail the novel method, Kernel Density Estimation Mixture Modelling (KDEMM). Finally, the two datasets used in this work are described, the synthetic data with known ground truth, used to compare models, and cognitive test data collected from the Dementia Research Centre (DRC), London, to derive the first event-based models of cognitive deterioration in PCA and tAD.

### A. Event-Based Model

The Event-Based Model (EBM) (Fonteijn et al. 2011) is used to estimate the ordering of biomarkers which move outside of a normal healthy range in a population as a result of a disease. Previous formulations of the EBM have been used to estimate the order of brain region volume loss in sporadic Alzheimers (Young et al. 2014), familial Alzheimers and Huntingtons disease (Fonteijn et al. 2012). In these versions of the EBM, parametric mixture modelling is used to predict the probability of an event having occurred, which the term given to a measurement transitioning from a healthy biomarker range to one associated with a disease.

In the EBM two component univariate mixture models are fit for each individual biomarker. The probability of belonging to each component of a mixture model is then used to as the probability of an event having occurred, *P*(*x_ij_|*E*_i_*), and an event not having occurred, *P*(*x_ij_|-E_i_*). These probabilities are used to give the likelihood of an ordering of biomarkers, *S*, using

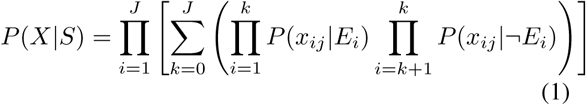

where *P*(*x*_*ij*_|*E*_*i*_) and *P*(*x_ij_|-E_i_*) are the probability of a measurement *x* ∈ *X* given an event having occurred and not occurred respectively, *i* ∈ *I* is the biomarker index and *j* ∈ *J* is the participant number.

When only a small number of biomarkers are used, all possible orderings can be enumerated and the characteristic ordering, 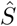, is the sequence which maximises *P*(*X|S*) (Equation (1)). As the number of possible orderings grows factorially as the number of biomarkers increases, Markov chain Monte Carlo (MCMC) sampling is used to find 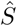 when the number of sequences to sample is too large.

Once we have 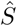, it can be used to estimate a disease stage for individuals given their biomarker measurements. The disease stage, *k*, is defined as, the stage, i.e. the number of events that have occurred, that has the highest probability given the data and our sequence, this is calculated using Equation (2).

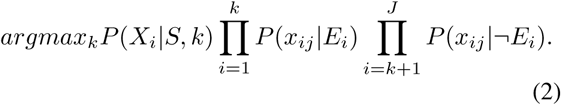

### B. Mixture Modelling

The steps detailed in Section II-A require models for *P*(*x*_*ij*_|*E*_*i*_) and *P*(*x*_*ij*_|-*E*_*i*_). Similarly to previous work using the EBM, here we will use two component Gaussian Mixture Models as our GMM models.

Initial parameters and constraints for each GMM component were derived from labelled data by taking the mean and standard deviation of each subpopulation, the mixture coefficient was initialised to 0.5 and constrained to the range [0.1, 0.9]. Parameters were then optimised to minimise the negative log-likelihood of the data given the model, using the Sequential Least SQuares Programming (SLSQP) algorithm. Constraints, initial parameters and the SLSQP algorithm were chosen similarly to previous implementations of GMM’s in disease progression modeling (Young et al. 2014; Fonteijn et al. 2011; Fonteijn et al. 2012).

### C. Kernel Density Estimation

Kernel Density Estimation (KDE) is a non-parametric method of probability density estimation, that is useful for data smoothing. The KDE estimation, 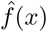, of a are the probability of function, *f*(*x*), with a independent and identically distributed sample, (*x*_1_*, x*_2_, …, *x*_*n*_), drawn from a distribution with an unknown density, is given by

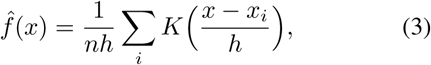

where *K* is non-negative function which integrates to one and has mean zero, and *h* is a positive smoothing factor called a bandwidth. With an appropriate choice of K, KDE naturally extends to multivariate density estimation.

In this work we use the scikit-learn (Pedregosa et al. 2011) implementation of KDE, using default parameters, including Gaussian kernel, for all values except the bandwidth, which was estimated using Scotts normal reference rule (Scott 1979).

### D. Kernel Density Estimation Mixture Modelling

To allow accurate fitting of mixture models with unknown underlying distributions we have implemented a novel algorithms for non-parametric mixture modelling. Let 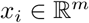 be a set of observations, and 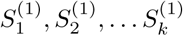 be known subsets of the data. The bandwidth *h* of these data is then estimated by applying Scotts rule to all the observations. Mixture weights 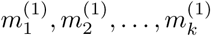 are initiated as 1/*k* for all subsets. Similarly to the *k*-means algorithm, the KDEMM algorithm then iterates over alternating assignment and update steps to optimise parameters (Figure 1).

#### Update Step

For each subset of the data, 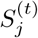, a KDE mixture component, 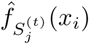, is fit using

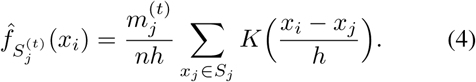

#### Assignment Step

Each observation is then assigned to a new subset, 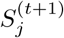, to which it has the maximum likelihood of belonging, i.e. 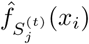. Mixture weights are then updated to be the proportion of observations in each subset i.e. 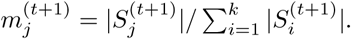.

The update and assignment steps are then iterated until subset assignment is no longer updated, i.e 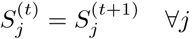. In this work an additional constraint, 0.1 < *m*_*j*_ < 0.9 ∀_*j*_, is placed on the mixture weights to ensure that subsets do not vanish, similarly to the GMM.

### E. Simulated Data

Since the ground truth is not known for most real-world problems, we chose to synthesise biomarker data to test the effect of using different mixture modelling techniques in the EBM framework. Two subpopulations were created: CN (*n* = 100) and AD (*n* = 100), corresponding to healthy controls and individuals with a disease, respectively. For each individual three synthetic biomarker measurements were generated, with each biomarker having two possible distributions corresponding an event having occurred and not occurred respectively (Figure 2). For each dataset a randomly generated event-sequence was generated, and the progression of biomarkers from normal to abnormal followed this event sequence. For the CN subpopulation 55% were assigned to stage zero (no biomarker events occurred), 25% at stage one (the first biomarker event has occurred and others have not), 15% at stage two (first two biomarker events have occurred) and 5% at stage three (all biomarker event have occurred). For the AD subpopulation, this trend is reversed with 55% at stage three, 25% at stage two, 15% at stage one and 5% at stage zero.

In this work we used Gamma distributions with varying parameters to synthesise biomarker data. The Gamma distribution was chosen because the shape parameter can be adjusted to yield distributions with different shape profiles. Using the scipy library (Jones et al. 2001), two sets of Gamma-distributed random numbers were generated, using default scale and location parameters. Both sets were mean-centred and one was mirrored by multiplying by negative one. The factor *f* = 2 × *ϕ* × *σ*_*k*_ is the ratio of standard deviation that separate the two components, where *ϕ* is a variable altered in each experiment and *σ*_*k*_ is the standard deviation of a Gamma distribution with the shape parameter *k*. The sets were then separated by adding *f* to one of the sets. This separation factor was used to simulate varying levels of separation between groups that is observed in cognitive test results. For each shape parameter and separation factor *n* = 25 different datasets were generated (Figure 2).

To compare models using simulated data we used three measures: likelihood of the data given the mixture model; correlation of predicted sequence with the ground truth; and correlation of EBM stage to the ground truth stage. Likelihood is used as a goodness of fit model, to test which of the mixture models explains the data better. To compare predicted sequences with the ground truth we use the Kendall-Tau rank correlation coefficient, which is a non-parametric test of how similar two sequences are, with *τ* = −1 implies that one sequence is the mirror opposite of the other and *τ* = 1 implies that the two sequences are identical. Finally we use the Spearman’s rank correlation coefficient to compare the accuracy of the maximum-likelihood stage (Section II-A) with the ground truth stage used to generate the data.

### F. Patient Data

Individuals with a clinical diagnosis of PCA and tAD were recruited between October 2005 and June 2016 at the Dementia Research Centre, London. 81 participants with PCA, 61 participants with tAD and 23 controls from the Young Onset Alzheimer’s Disease (YOAD) study (Table I). Patients attended the Cognitive Disorder clinic at the National Hospital of Neurology and Neurology, or were recruited by individual referral from other neurologists to whom they expressed interest for taking part in observational research. All PCA patients met both Tang-Wei (Tang-Wai et al. 2004) and Mendez (Mendez, Ghajarania, and Perryman 2002) criteria based on available information at baseline and expert retrospective clinical review. Participants were excluded if they also met criteria for another neurodegenerative syndrome, thus fulfilling consensus criteria for PCA-pure (Crutch et al. 2017). Patients with PCA and patients with typical Alzheimer’s disease fulfilled research criteria for probable Alzheimer’s disease (Dubois et al. 2010; Dubois et al. 2007).

**Fig. 1:**
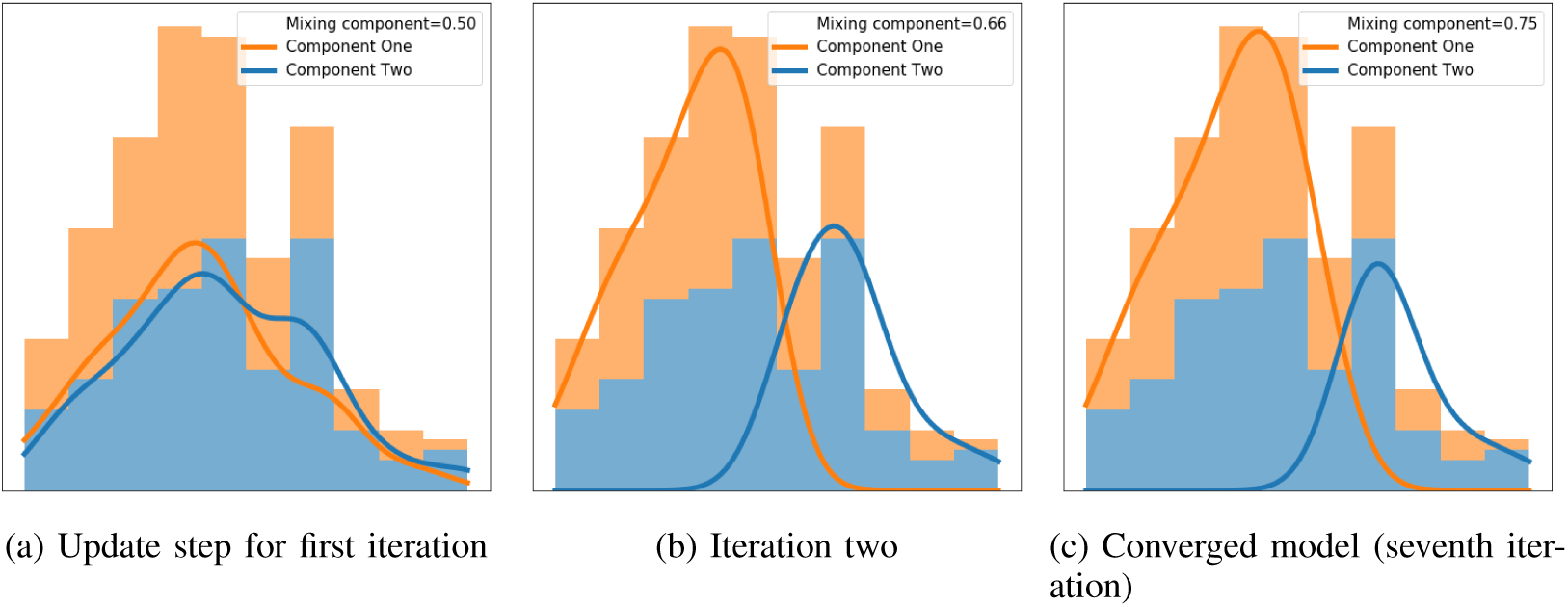
Example fitting process for a KDEMM.

**Fig. 2:**
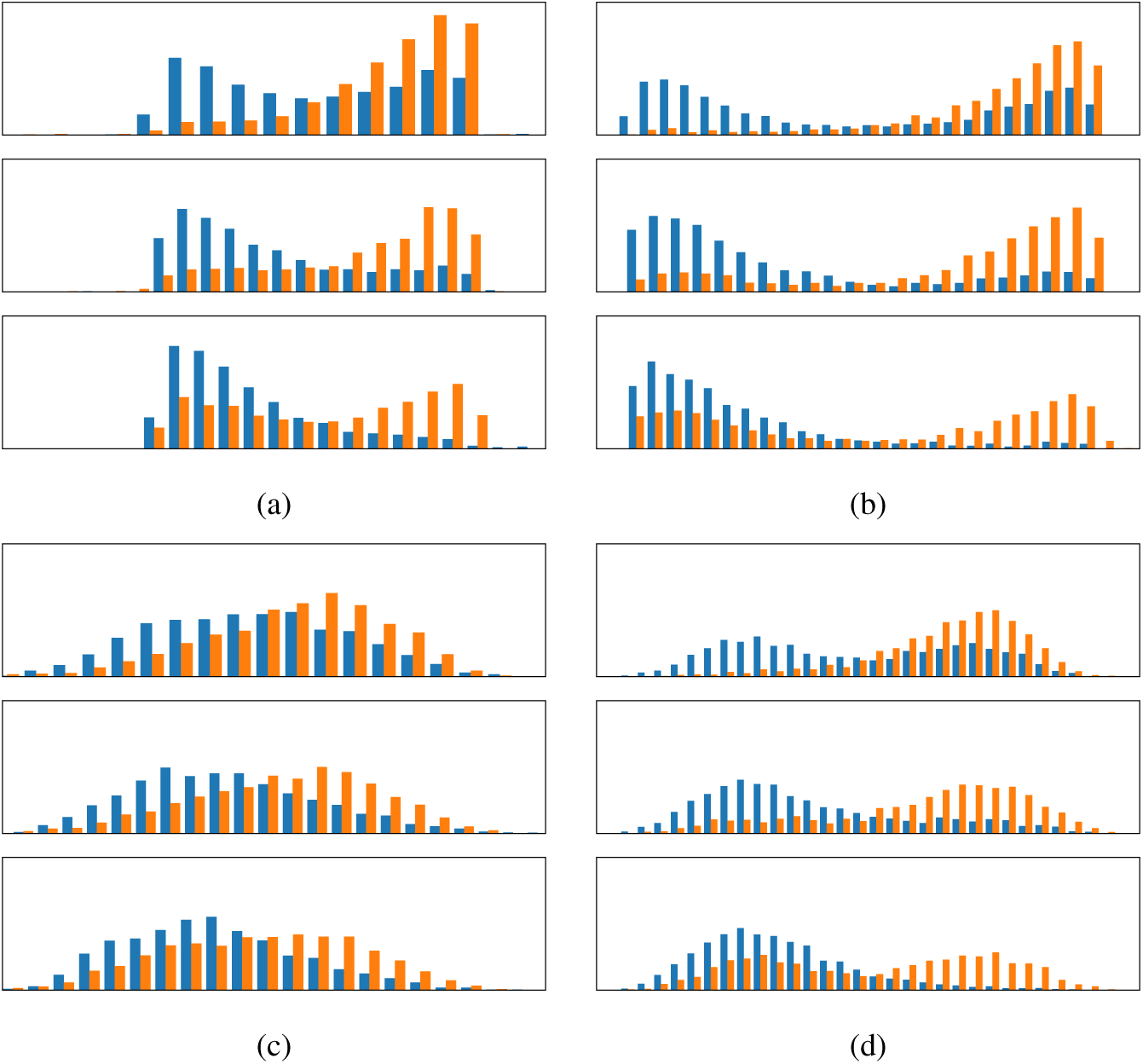
Histograms of exemplar synthetic data. Each dataset is comprised of 5000 subjects, each with three biomarker measurements, with disease sequence 1, 2, 3. a) and b) have a shape parameter *k* = 1.5 and c) and d) have *k* = 10. a) has a separation factor of *sf* = 1, b) *sf* = 2, c) *sf* = 0.5, d) *sf* = 1.5.

To compare the performance of the GMM and KDEMM on this cognitive dataset, both models were used to fit EBMs for the baseline visits from both the PCA and tAD subgroups, resulting in Maximum Likelihood (ML) sequences of events for both. To measure the confidence in these ML sequences, 100 models were fit on bootstrap resampled datasets, for both models and PCA and tAD datasets. Bootstrapping was performed by randomly sampling, with replacement, dataset of the same size as the original samples,(PCA= 81, tAD= 61) and both mixture models and maximum likelihood sequences were fit for each of these 100 random samples. We chose to use the same fitting procedure as previous EBM literature (Oxtoby et al. 2017; Young et al. 2014) to isolate the contribution of the KDEMM.

**TABLE I:**
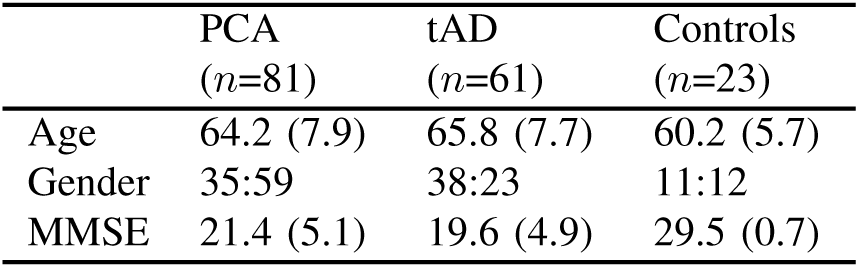
Demographics of participants.

## III. Results

In this section GMMs and KDEMM are compared, firstly by examining the likelihood of the data given each mixture model. This comparison shows which technique is most suitable for modelling the data as a two-component mixture. Secondly, these models are compared in the context of the EBM in two ways: ability to recreate the event sequence of simulated data and also the ability to accurately stage simulated data. These two comparisons are used to test the hypothesis that the KDEMM is more appropriate in the EBM for non-Gaussian data.

### A. Mixture model comparison

For each parameter combination (Section II-E), *ϕ* (*n* = 8) and *k* (*n* = 9), 25 datasets consisting of three biomarkers were synthesised as described. For each of the 5,400 (8 × 9 × 25 × 3) biomarkers, both a GMM and KDEMM were fit, and the likelihood of the data under the models was calculated. Figure 3 shows that the GMM has a higher likelihood than the KDE for most parameter combinations, particularly when the shape parameter of the Gamma distribution is small, i.e. the distribution more skewed. This likely is a result of the standard deviation of these datasets being very small, resulting in very high outputs from the probability density functions from the fit Gaussian distributions. The negative log likelihood of the data given the model was significantly lower for the GMM model (355.69) compared to the KDE (370.40, *p* < 1*e* − 14) model across all the trials.

As the intended use for these mixture models is to predict probability of an event having occurred in the EBM, the performance of both the GMM and KDEMM are compared in the EBM. Synthetic datasets were generated with random event sequences, and for each model the characteristic event sequence was generated by enumerating all possible sequences. Figure 4 shows the difference between the GMM and KDEMM EBMs in Kendall-Tau correlation coefficient with ground truth. It can be seen that the EBM using the KDEMM has a significantly higher correlation (*τ* = 0.89) with the ground truth order of events compared to EBM using GMM (*τ* = 0.50, *p* < 1e − 90). This trend is seen across the majority of parameters used to create the synthetic datasets. However the GMM performs equally well when there is a sufficiently large separation between components and large enough shape parameter (thus reducing the skewness, i.e. more similar to Gaussian). The GMM performs better in only three parameter combinations, and in only one of these, *ϕ* = 10 and *k* = 10, does the corresponding EBM perform significantly better.

**Fig. 3:**
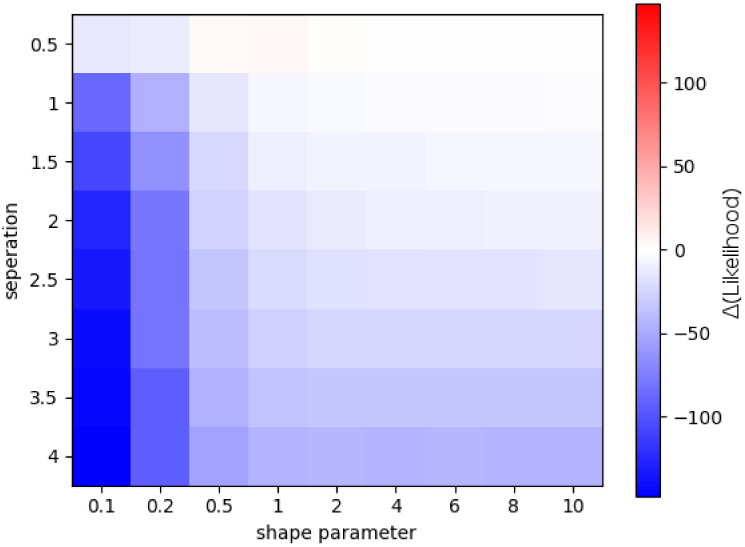
Heat map showing difference in likelihood of data between a KDEMM and GMM. Red indicates that KDEMM fits the data better and blue indicates that the GMM fits the data better.

As well as using the EBM for generating an event sequence, it has also been used to give a disease stage for study participants. Figure 5 shows a comparison between modelled stages from GMM- and KDEMM-EBMs built using the ground truth stage. It can be seen that the EBM stages using the KDEMM model, correlated with the ground truth stages (*ρ* = 0.88) significantly better than the EBM stages using the GMM model (*ρ* = 0.83, 1e −18). Similarly to the event sequence the GMM staging performs equally well when the data was generated with a sufficiently large separation factor and shape parameter. This likely is due to the staging reliance on an accurate event order as well as well-fit mixture models.

**Fig. 4:**
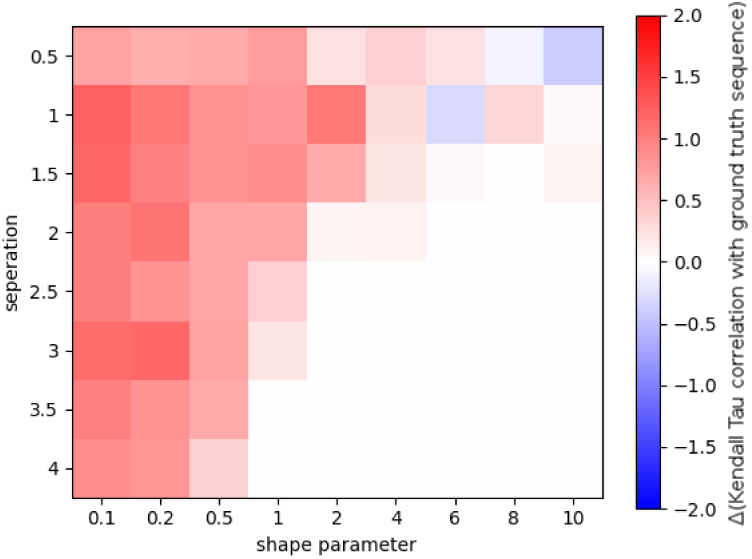
Heat map showing difference between the Kentall-Tau correlation with a randomly geneated sequence and the sequence derived from GMM and KDEMM. Red indicates that the KDEMM derived sequence correlates better and blue indicates that the GMM derived sequence correlates better.

**Fig. 5:**
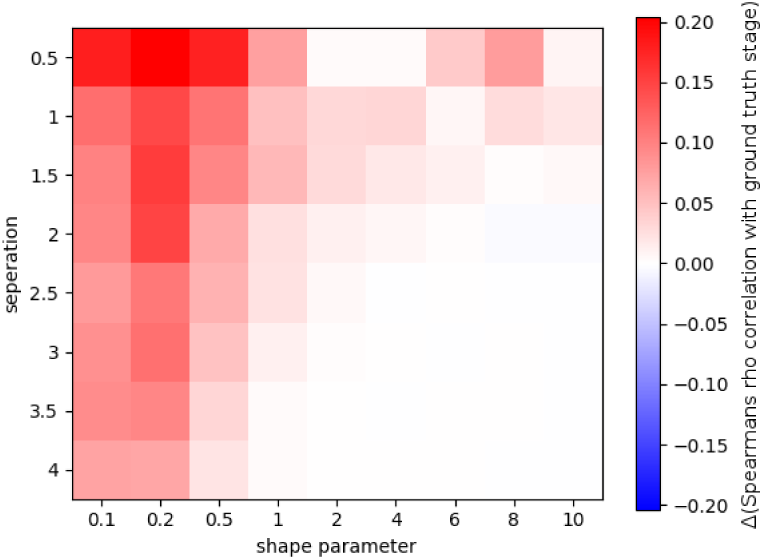
Heat map showing difference between the Spearman *ρ* correlation with a synthetic patients stage and the stage predicted by EBMs and NPEBMs. Red indicates that the NPEBM sequence correlates better and blue indicates that the EBM sequence correlates better.

### B. Application to cognition in Alzheimer’s

Using both the GMM and KDEMM models within the EBM for cognitive data, we see different distinct event orderings from each model. In the PCA Maximum Likelihood (ML) sequence fit using GMM models the Digit Span (F) and Digit Span (F Max.) are separated by 13 positions in the event sequence (Figure 6a); this is contrary to previous results (Lehmann et al. 2012), which would suggest that these subscores of the same test would occur at very similar positions in the event sequence. The ordering of cognitive tasks also does not fit with clinical expectations, with PCA patients showing earlier memory change (mean 33.2 ± 2.6 positions earlier) compared to tAD and no difference in visual change (mean 0.4 ± 5.9 positions earlier) compared to tAD patients. By comparison, ML sequences fit for both tAD and PCA using the KDEMM model yielded successive positions for the highly related Digit Span (F) and Digit Span (F Max) scores. As well as being clustered next to each other in the sequences, high uncertainty about the relative position of these tests can be observed, in both the direct fit (Figures 7a and 7c) and bootstrapped samples (Figures 7b and 7d), as the model is unable to accurately predict which comes first. The order of cognitive events using the KDEMM model (Fig 7) align much better than those using the GMM model with clinical definitions of these two conditions. Change on the five principle visual tests (A cancellation time, fragmented letters, dot counting, shape discrimination, object decision) was observed earlier in PCA than tAD in all cases (mean 5.6 ± 4.2 positions earlier), whilst position values for the three principle episodic memory tasks (short Recognition Memory Test [sRMT] for words and faces, Paired Associate Learning test [PAL]) were equivalent or earlier in tAD than PCA (mean 4.3 ± 5.9 positions earlier). In both the GMM- and KDEMM-EBMs there is a notable amount of uncertainty in the bootstrapped sequences, however in the GMM-EBM this appears to be distributed away from the ML sequence, as observed by the spread away from the diagonal (Figures 6b and 6d), whereas in the KDEMM-EBM the uncertainty is more focused on the diagonal (Figures 7b and 7d), suggesting that the sequence is more robust to bootstrapping. The broad uncertainty in the bootstrapped GMM-EBMs is likely due to the distributions being sampled during bootstrapping not being suitable for GMMs.

Assigning the ML stage for each participant’s baseline visit, the EBM is able to estimate the stage of the disease that each participant is at. Figure 8 shows the stages assigned by the tAD and PCA sequences in both GMM- and KDEMM-EBMs. It can be seen in across all the staged data that controls are generally assigned a low stage, and tAD and PCA participants are assigned higher stages, with the majority at the later stages of the disease (Figure 8). Notable exceptions are a small number of controls assigned to later stages in both GMM-EBMs (Figures 8a and 8c) and a small number of tAD participants who have been assigned early stages in the KDEMM model (Figure 8d). To analyse the consistency of the fitted models for PCA and tAD we staged each of the subsequent visits for all of the participants. These data were not used to fit the EBM, so provide a suitable test set. The PCA sequence from the GMM-EBM staged 106 of the 118 follow ups higher or the same as the baseline visit and 186 follow up visits staged higher or the same as a previous visit (201 comparisons made). For the KDEMM-EBM these numbers were 113 and 187 respectively. For the tAD EBM fit using GMMs 31 of the 32 follow ups were staged higher or the same as baseline visit and 36 visits staged higher or the same as a previous visit (37 comparisons made), for the EBM fit using KDEMMs these numbers were 26 and 31 respectively.

**Fig. 6:**
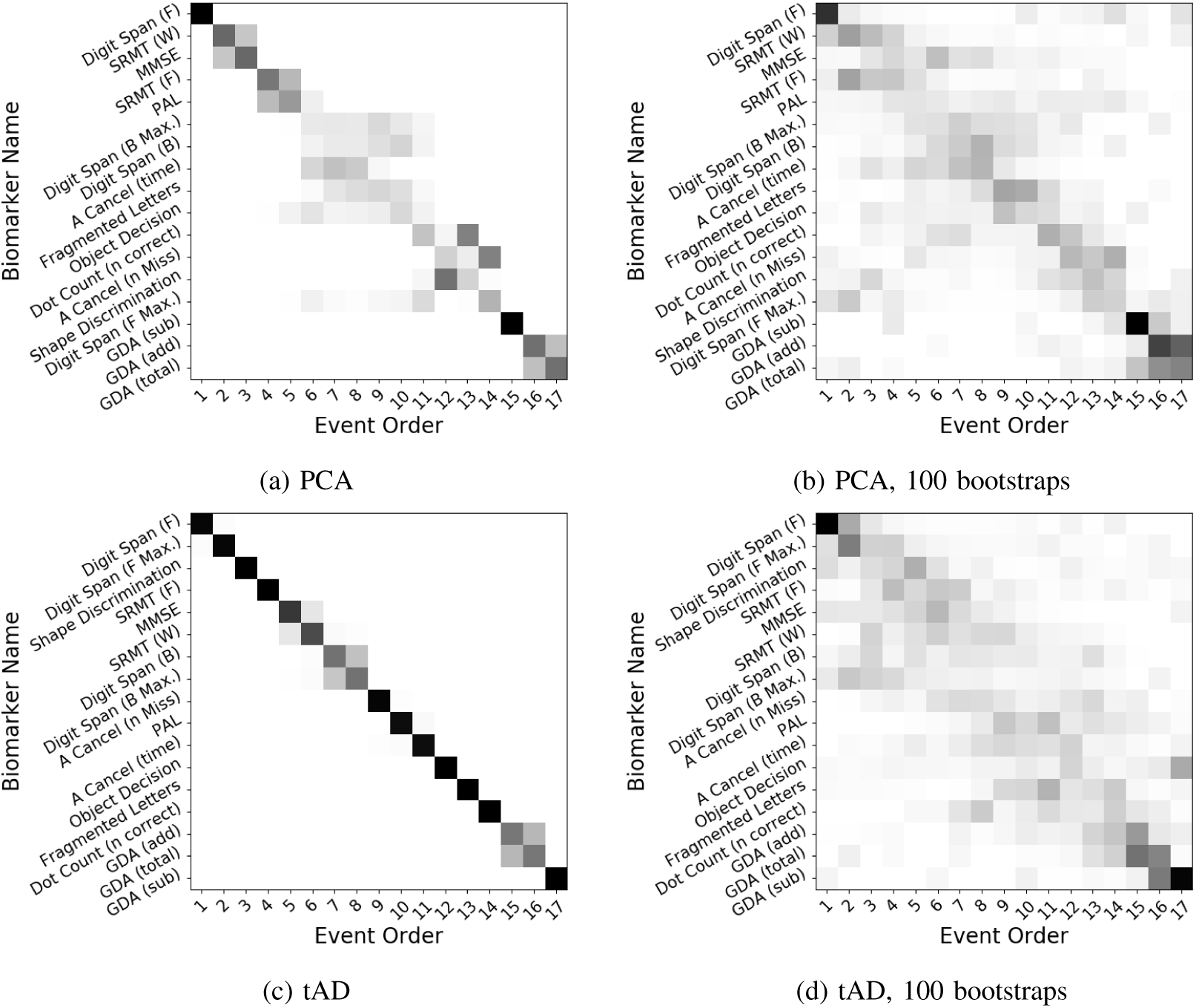
Positional variance diagram for EBM, showing uncertainity in the maximum likelihood sequence. Uncertaintity is measure by boostrapped resampling of the data 100 times, fitting an EBM on these bootstraps and plotting the positional variance of the Markov Chain Monte Carlo (MCMC) samples. Each entry in the positional variance diagram represents the proportion of the bootstrapped MCMC samples in which events appear at a particular position in the ML sequence (*x*-axis). This proportion ranges from 0 in white to 1 in black. The *y*-axis orders events by the maximum likelihood sequence.

## IV. Discussion and Conclusion

In this work we have introduced Kernel Density Estimation Mixture Models (KDEMM), a novel semi-supervised clustering algorithm, which we have used to adapt the Event-Based Model (EBM) to handle non-Gaussian-distributed data, in particular focusing on cognitive test data. We compared the KDEMM with the current state of the art mixture model technique used in the EBM. We compared mixture modelling techniques using both synthetic data and also cognitive test data in tAD and PCA. To understand isolated model performance we calculated the likelihood of the data for both GMM and KDEMMs across all datasets. These likelihoods suggested that the GMM is better at modelling the data, however this particular metric may not be suitable for an unbiased comparison of these models. As the GMM parameters were estimated by minimising the negative log likelihood of the data, and the KDEMMs were optimised to have stable clusters, it is unsurprising that the GMMs have a higher likelihood. Perhaps a more robust comparison of the two models would be to compare the ability to model underlying parameters used to generate data, however this is outside the scope of the current study as the EBM does not require this information.

**Fig. 7:**
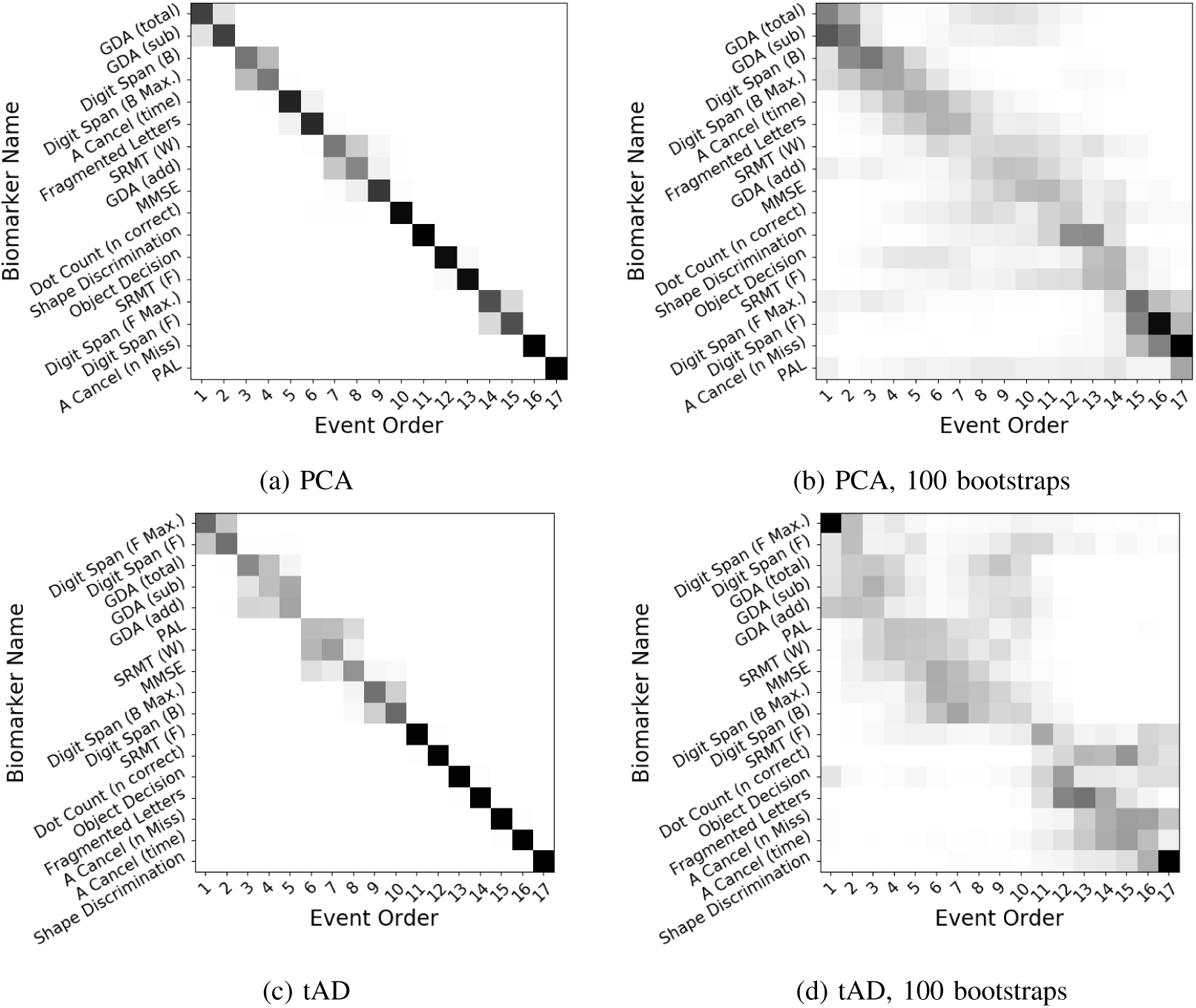
Positional variance diagram for NPEBM showing uncertainity in the maximum likelihood sequence. Uncertaintity is measure by boostrapped resampling of the data 100 times, fitting an NPEBM on these bootstraps and plotting the positional variance of the Markov Chain Monte Carlo (MCMC) samples. Each entry in the positional variance diagram represents the proportion of the bootstrapped MCMC samples in which events appear at a particular position in the ML sequence (*x*-axis). This proportion ranges from 0 in white to 1 in black. The *y*-axis orders events by the maximum likelihood sequence.

Results show that the using the KDEMM in the EBM yielded either more or equally accurate event sequences compared to the GMM, for the majority of parameters tested. The KDEMM performed particularly well on distributions that are more skewed and less separated, suggesting that, as intended, the KDEMM is better at modelling non-Gaussian data compared to the GMM. This result is in contrast with the previous result that showed that GMM models resulted in better fit models as measured by the data likelihood. As well as performing better on more skewed data the KDEMM also performed no worse than the GMM on almost all of the datasets tested, suggesting that it might be a more suitable choice for all data types. Using the event sequences to stage all synthetic data showed that the KDEMM model was significantly better than the GMM, however the magnitude of difference is quite small and as the staging is reliant on the event-order this result is not surprising.

We applied both mixture modelling techniques in an EBM to a dataset of neuropsychological test results in PCA and tAD, different AD phenotypes that are predominantly led by visual cognitive impairment or memory respectively. The KDEMM provided an event ordering which was more in line with clinical experience in both PCA and tAD compared to the GMM, with impairment on visual tasks seen earlier in PCA and impairment on episodic memory tasks seen earlier in tAD. The KDEMM also demonstrated more certainty in the maximum likelihood sequences, as estimated by bootstrapping, which is itself prone to over estimating error. Using the fitted models to estimate disease stages of follow up data for people with PCA and tAD, showed that the KDEMMM-EBM provided a more monotonically progressive model in PCA compared to the GMM-EBM, however this trend was reversed when looking at the tAD population. Closer inspection of the four tAD participants with decreasing disease stages showed that learning effects were seen in nine of the 17 tests, making the results of this comparison difficult to interpret. A larger number of tAD follow ups would be required to reach a more definitive conclusion about the disease staging.

**Fig. 8:**
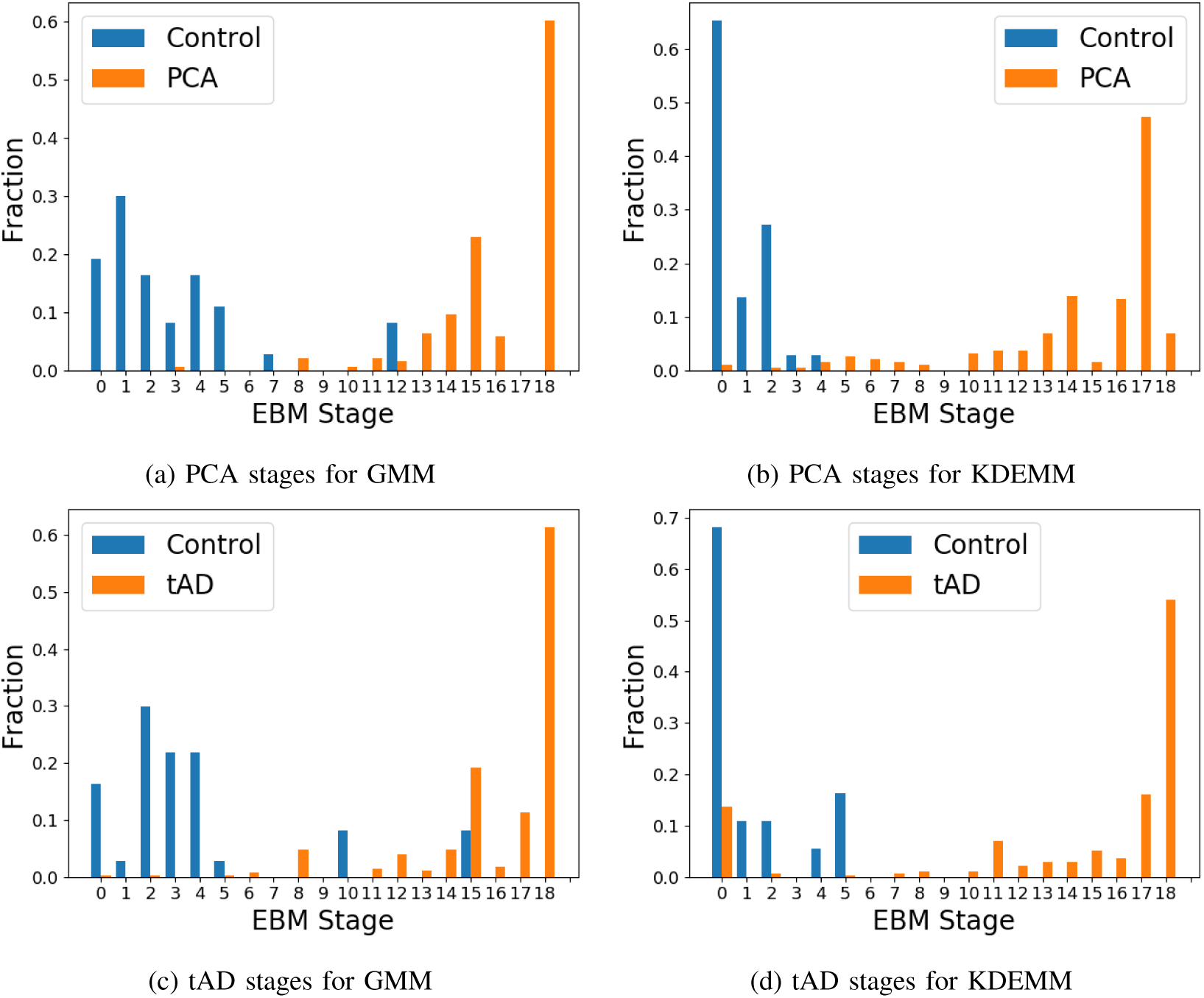
Histogram of the maximum likelihood stages assigned to controls.

As with most previous descriptions of the EBM the key limitation to the model is that it assumes a single event sequence, whereas clinical data would suggest that both tAD and PCA are heterogeneous diseases which have many possible patterns of evolution. Though recent work in this area has adapted the EBM framework to enable multiple event sequences (Young et al. 2017). A limitation of the KDEMM compared to the GMM it its reliance on data, as Kernel Density Estimation utilises all data to make a prediction, compared to the two parameters in the Gaussian distribution. Though both methods would perform worse with less data, it is possible in incorporate prior knowledge to parametrise a GMM, for example normative data from cognitive tests, whereas this is not possible using a KDEMM. Another limitation of the KDEMM is the computational complexity to both fit and apply the models on large datasets; it requires considerably more time to fit than the GMM.

The KDEMM extends to a number of potential applications outside of the EBM framework. Future work will include the use of KDEMM outside of this framework in applications such as disease subtype clustering and feature normalisation for supervised learning. Though already used in two studies (Oxtoby et al. 2017; Firth et al. 2018), future work includes applying the KDEMM EBM to other multimodal datasets and comparing it to results from the GMM. Finally, as well as using the classic EBM approach, future work will also include integrating the KDEMM into the recently proposed Discriminative EBM (Venkatraghavan et al. 2017), to test whether our approach makes improvements to this method.

## Ethics

This project was approved by the NRES Committee London, Queen Square, and all participants provided written informed consent according to guidelines established by the Declaration of Helsinki.

## Funding

This work was supported by EPSRC (EP/M006093/1). The cognitive data collection was supported by grants to SC from ESRC/NIHR (ES/L001810/1) and Alzheimers Research UK (Senior Research Fellowship). NPO and DCA acknowledge that their contributions to this work is part of a project that has received funding from the European Unions Horizon 2020 research and innovation programme under grant agreement number 666992. ALY is supported by an EPSRC doctoral prize fellowship.

## Conflict of Interest Statement

The authors declare that the research was conducted in the absence of any commercial or financial relationships that could be construed as a potential conflict of interest.

## ACKNOWLEDGEMENTS

We would like to thank all participants in the PCA longitudinal study for donating their time to research. We would also like to the thank everyone at the Dementia Research Centre who helped with the data collection over the course of twenty years, including Manja Lehmann, Tim Shakespeare, Aida Suarez Gonzalez, Amelia Carton, Ivanna Pavisic and Dilek Ocal. The Dementia Research Centre (Institute of Neurology, University College London, UK) is an Alzheimers Research UK Coordinating Centre and is grateful for support from the Department of Healths NIHR Biomedical Research Centres funding scheme, the NIHR Queen Square Dementia Biomedical Research Unit, the MRC Dementias Platform UK, and the Leonard Wolfson Experimental Neurology Centre.

